# Regulation of ATR activity by the RNA polymerase II phosphatase PNUTS-PP1

**DOI:** 10.1101/267013

**Authors:** Helga B. Landsverk, Lise E. Sandquist, Gro Elise Rødland, Beata Grallert, Laura Trinkle-Mulcahy, Randi G. Syljuåsen

## Abstract

Ataxia telangiectasia mutated and Rad3-related (ATR) kinase is a key factor activated by DNA damage and replication stress. Here, we show that ATR signaling is increased in human cells after depletion of the RNAPII phosphatase PNUTS-PP1, which dephosphorylates RNAPII on Ser 5 of its carboxy-terminal domain (CTD) (pRNAPII S5). Increased ATR signaling was observed in the presence and absence of ionizing radiation or replication stress and even in G1 phase after depletion of PNUTS. Vice versa, ATR signaling was reduced, in a PNUTS dependent manner, after inhibition of the CDK7 kinase mediating pRNAPII S5. Furthermore, CDC73, a well-known RNAPII-CTD binding protein, was required for the high ATR signaling after depletion of PNUTS and co-immunoprecipitated with RNAPII and ATR. These results suggest a novel pathway involving RNAPII, PNUTS-PP1 and CDC73 in ATR signaling and give new insight into the diverse functions of ATR.

## Introduction

The ataxia telangiectasia mutated and Rad3-related (ATR) kinase is a master regulator of DNA damage and replication stress signaling coordinating DNA repair, cell cycle checkpoint and cell death pathways (Cimprich & Cortez, 2008). Understanding how ATR is activated is therefore a critical issue in biomedical research. The canonical pathway for ATR activation is initiated by the presence of single stranded DNA (ssDNA) coated by RPA (ssDNA-RPA) (Zou & Elledge, 2003). ssDNA-RPA at sites of DNA damage recruits ATR via its obligate binding partner ATRIP (Cortez, Guntuku et al., 2001, Zou & Elledge, 2003). Full activation of ATR is further facilitated by TOPBP1 (Cimprich & Cortez, 2008). A large amount of evidence supports an important role for the canonical pathway in ATR activation (e.g. reviewed in (Marechal & Zou, 2015)) However there is also evidence suggesting the existence of alternative pathways (Nam & Cortez, 2011), which are less well understood.

One tentative alternative pathway proposes that the cell takes advantage of its transcription machinery to activate ATR (Derheimer, O’Hagan et al., 2007, Lindsey-Boltz & Sancar, 2007). This was proposed based on the finding that stalled, elongating RNAPII could induce ATR-dependent P53 phosphorylation in the absence of DNA damage (Derheimer et al., 2007). RNAPII is a recognized sensor in transcription coupled repair where it recruits DNA repair factors to sites of damage (Andrade-Lima, Veloso et al., 2015, Spivak, 2016). The discovery of pervasive transcription outside protein coding genes (Jensen, Jacquier et al., 2013), suggests RNAPII might be scanning a majority of the genome and makes an involvement of RNAPII in sensing DNA damage and activating ATR more likely (Lindsey-Boltz & Sancar, 2007). However, such an upstream role of RNAPII in ATR activation has yet to gain wide acceptance, perhaps because the factors involved in signaling between stalled RNAPII and ATR remain unknown.

During the transcription cycle, RNAPII becomes reversibly phosphorylated on the carboxy-terminal domain (CTD) of its largest subunit. Phosphorylation of specific residues in the CTD heptapeptide repeats, ie Ser 2 (S2) and Ser 5 (S5), is associated with specific phases of the transcription cycle. This is thought to contribute to a CTD ‘code’, in which combinations of post-translational modifications on the CTD can be ‘written’ and ‘read’ to regulate association with transcription and RNA processing factors (Egloff, Dienstbier et al., 2012). Interestingly, increased phosphorylation of the CTD has been observed after ultraviolet radiation and camptothecin in human cells (Rockx, Mason et al., 2000, Sordet, Larochelle et al., 2008) and is tightly connected to RNAPII stalling (Alexander, Innocente et al., 2010, Boehm, Saunders et al., 2003). Notably, RNAPII stalling can also occur after other types of DNA damage, e.g. following ssDNA breaks or cyclopurines such as formed after IR (Andrade-Lima et al., 2015, Brooks, Wise et al., 2000, Cadet, Davies et al., 2017, Kathe, Shen et al., 2004). Furthermore, several proteins that interact with the phosphorylated CTD were required for resistance to ionizing radiation (IR) or doxorubicin in *Saccharomyces cerevisiae* (Winsor, Bartkowiak et al., 2013). Based on these findings, one possibility would therefore be that RNAPII responds to DNA damage by signaling via its CTD.

We previously discovered that the <u>P</u>rotein Phosphatase 1 <u>N</u>uclear <u>T</u>argeting <u>S</u>ubunit (PNUTS) suppresses the G2 checkpoint after IR, but the underlying molecular mechanisms remained to be identified (Landsverk, Mora-Bermudez et al., 2010). Interestingly, PNUTS is one of the most abundant nuclear regulatory subunits of PP1 (Jagiello, Beullens et al., 1995, Kreivi, Trinkle-Mulcahy et al., 1997), and RNAPII CTD is the only identified substrate of PNUTS-PP1 (Ciurciu, Duncalf et al., 2013). PNUTS-PP1 dephosphorylates RNAPII S5 (CTD) in vitro (Lee, You et al., 2010) and depletion of PNUTS causes enhanced RNAPII S5 phosphorylation (pRNAPII S5) in human cells (Xing, Lin et al., 2014). Because RNAPII, as described above, has a proposed role in ATR activation and ATR is a crucial player in the G2 checkpoint, we addressed whether PNUTS-PP1 might suppress ATR signaling. Our results show that ATR signaling increases after PNUTS depletion in a manner not simply correlating with DNA damage or replication stress. The increased ATR signaling rather appears to depend upon CTD phosphorylation, which is counteracted by PNUTS-PP1. Furthermore, the known CTD-binding protein, CDC73, is required for the high ATR signaling and co-immunoprecipitates with ATR.

## Results

In our previous work (Landsverk et al., 2010), we observed increased phosphorylation of CHK1 and RPA32 at late timepoints (2-24 hr) after IR in PNUTS depleted HeLa cells. As CHK1 and RPA32 are ATR targets (Reaper, Griffiths et al., 2011, Shiotani, Nguyen et al., 2013), we addressed whether ATR signaling was affected specifically. Indeed, depletion of PNUTS with two different siRNA oligos caused increased IR-induced phosphorylation of the ATR substrates CHK1 S317 and RPA S33, but not of the ATM substrate CHK2 T68 (Fig 1A). Phosphorylation of CHK1 and RPA were increased both at early (5min-1h) and late (6h) timepoints after IR, as well as in the absence of IR (Fig 1A), suggesting a general role for PNUTS in suppressing ATR signaling. In agreement with this notion, pCHK1 S317 and pRPA S33 were higher also during thymidine-induced replication stress in PNUTS depleted cells (Fig 1B). Similar results were found in U2OS cells (Fig S1A), and the effect was clearly ATR-mediated, as the ATR inhibitor VE-821 inhibited the increased CHK1 phosphorylation after IR and thymidine (Fig S1B,C). Inhibition of ATR activity was not a general effect after depletion of a PP1 regulatory subunit because knockdown of another abundant nuclear regulatory subunit, NIPP1 (Jagiello et al., 1995), did not increase CHK1 S317 or RPA S33 phosphorylation (Fig S1D). Furthermore, the increased ATR signaling was not due to off-target effects of the siRNA oligonucleotides, since expression of mouse pnuts-EGFP to near endogenous levels abrogated the increased CHK1 phosphorylation after depletion of human PNUTS, both in the absence and presence of IR (Fig 1C).

**Figure 1.**
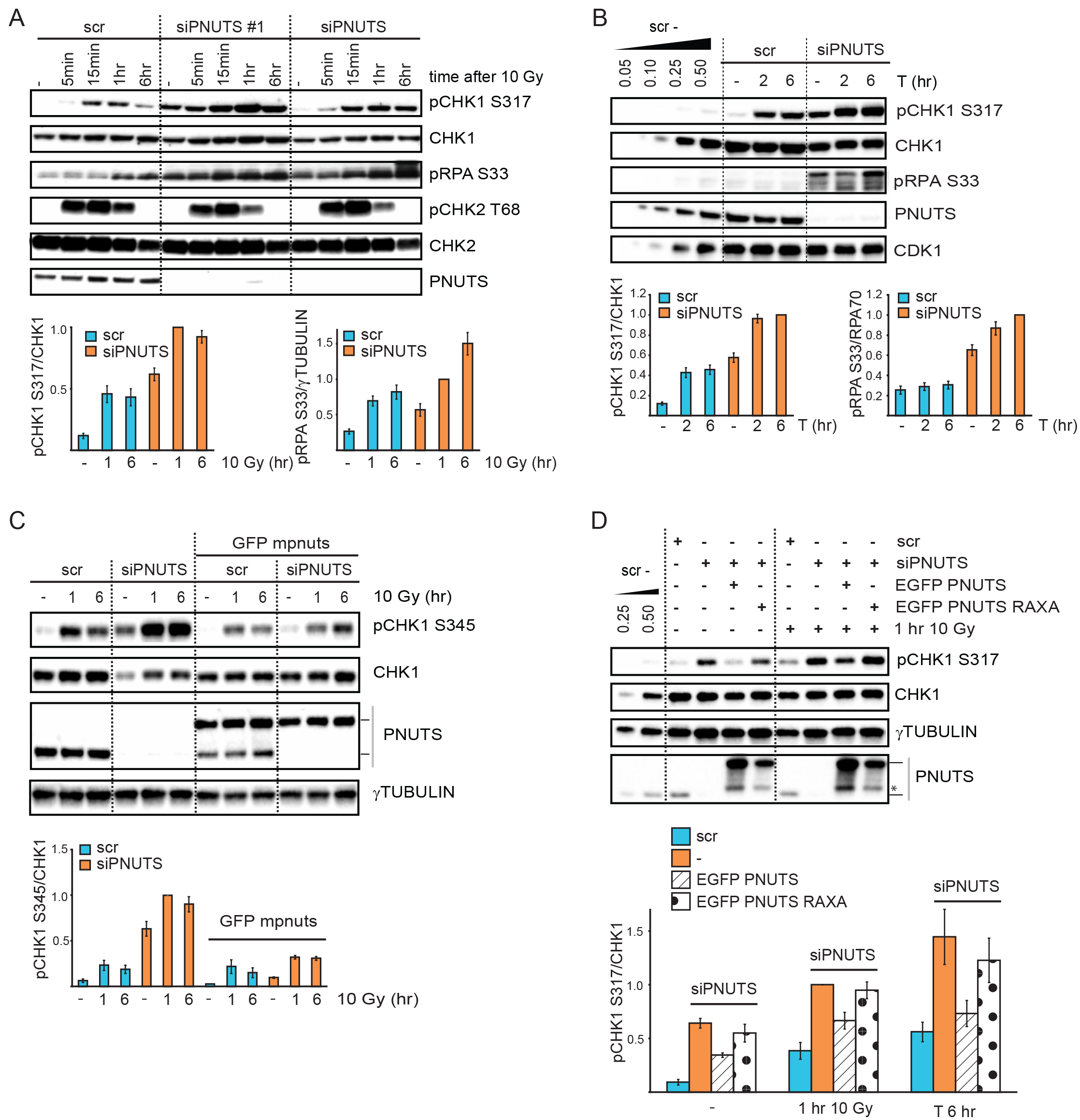
PNUTS-PP1 suppresses ATR signaling. **A)** Western blot analysis of ATR and ATM signaling events in control scrambled siRNA transfected (scr) or PNUTS siRNA transfected (siPNUTS #1 and siPNUTS) HeLa cells, at indicated times after 10 Gy. Cells were harvested at 72 hours after siRNA transfection. Bottom bar charts show quantification of pCHK1 S317 relative to CHK1 and pRPA S33 relative to γTUBULIN levels (n=8). **B)** Western blot analysis at 2 or 6 hr after addition of thymidine to cells siRNA transfected as in A) (scr and siPNUTS). Bottom bar charts show quantification of pCHK1 S317 relative to CHK1 and pRPA S33 relative to RPA70 levels (n=10). **C)** Western blot analysis of HeLa cells or HeLa BAC clones stably expressing EGFP mouse pnuts (mpnuts) transfected with scr or siPNUTS (specifically targets human PNUTS), at 1 or 6 hr after 10 Gy. Lines to the right of the western blot indicate migration of human endogenous PNUTS (lower band) and EGFP mpnuts (upper band). Bottom bar chart shows quantification of pCHK1 S345 relative to CHK1 levels (n=3). **D)** Western blot analysis of HeLa cells transfected with scr or siPNUTS. At 24 hr post transfection, the indicated samples were transfected with wild type EGFP PNUTS or PP1-binding deficient EGFP PNUTS RAXA. Cells were harvested 48 hr later without further treatment (-) or 1 hr after 10 Gy. Lines to the right of the western blot indicate migration of endogenous PNUTS (lower band) and EGFP PNUTS/EGFP PNUTS RAXA (upper band), asterisk indicates what is likely EGFP PNUTS/EGFP PNUTS RAXA degradation products. Bar chart shows quantification of pCHK1 S317 relative to CHK1 (n=3).

To address the importance of PP1 for the inhibitory effects of PNUTS on ATR signaling, siRNA-resistant wild type and PP1-binding deficient PNUTS were overexpressed in cells depleted of endogenous PNUTS. Wild type PNUTS, but not PP1-binding deficient PNUTS RAXA (Kreivi et al., 1997), partially abrogated increased CHK1 phosphorylation in the absence of exogenous stress and after IR or thymidine (Fig 1D and S2A), showing that PP1-PNUTS binding is important for the negative effect of PNUTS on ATR signaling. Higher expression levels of the PNUTS RAXA mutant did not alter these results (Fig S2B).

Potentially, PNUTS-PP1 could counteract ATR signaling by generally dephosphorylating ATR substrates, as is the case for *Saccharomyces cerevisae* PP4 and the ATR homologue Mec1 (Hustedt, Seeber et al., 2015). To address this, we added an ATR inhibitor after induction of ATR signaling by IR. If PNUTS-PP1 directly dephosphorylates CHK1 and RPA, depletion of PNUTS should cause delayed removal of pCHK1 S317/S345 and pRPA S33 after addition of the ATR inhibitor. However, both pCHK1 S317 and pRPA S33 declined at a similar rate in cells transfected with control siRNA and PNUTS siRNA (Fig 2A), showing that phosphatase activity against these substrates is similar under these conditions. Furthermore, overexpression of PNUTS did not decrease pCHK1 S317 or pRPA S33 (Fig 1D and data not shown). This suggests PNUTS-PP1 does not directly dephosphorylate these ATR targets. To further verify this finding, we also examined pCHK1 S317/S345 and pRPA S33 after addition of the ATR inhibitor to thymidine-treated cells transfected with control siRNA and PNUTS siRNA (Fig S2C). Decline of pCHK1 S317 and pCHK1 S345 occurred similarly also under these conditions, consistent with the notion that CHK1 is not a direct substrate of PNUTS-PP1. On the other hand, pRPA S33 declined less in PNUTS depleted cells in the presence of thymidine (Fig S2C). As pRPA S33 declined similarly in cells transfected with control and PNUTS siRNA after IR (Fig 2A), this most likely implies that another kinase contributes to pRPA S33 in PNUTS depleted cells after prolonged replication stress (thymidine 16h). ATR-independent phosphorylation of pRPA S33 has e.g. been reported in the presence of hydroxyurea (HU) in combination with ATR inhibitor (Toledo, Murga et al.). Altogether, these results nevertheless suggest that PNUTS-PP1 does not suppress ATR signaling by generally counteracting phosphorylation of its downstream substrates.

**Figure 2.**
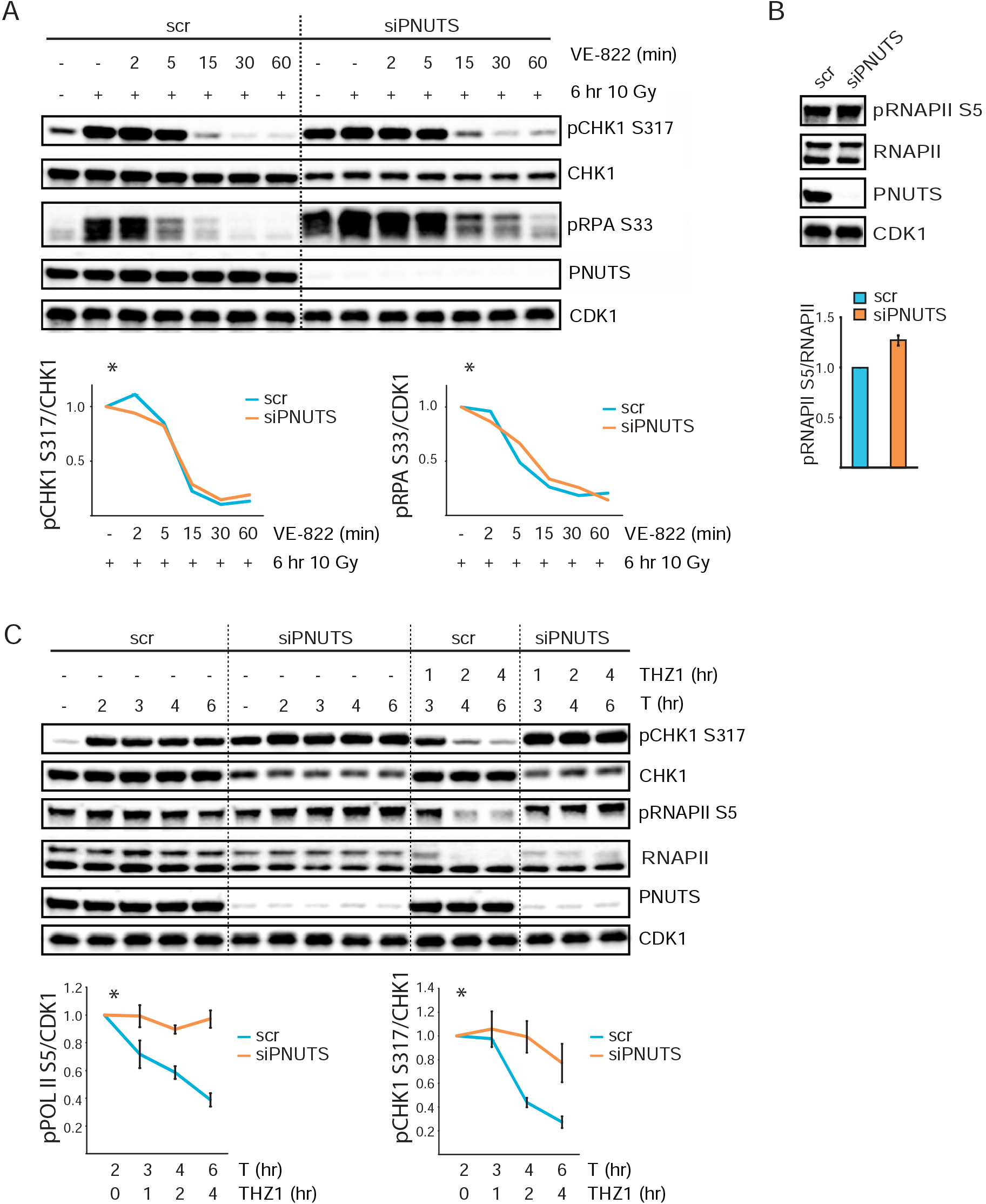
PNUTS-PP1 likely suppresses ATR signaling by dephosphorylating pRNAPII CTD. **A)** Western blot analysis of scr or siPNUTS transfected cells without IR or 6 hr after 10 Gy. VE822 was added for 2, 5, 15, 30, or 60 min to indicated samples 6 hr after 10 Gy. *Charts show values of pCHK1 S317/CHK1 in the VE822-treated cells rela-tive to 10 Gy 6hr, for respective siRNA oligos. Experiment was performed 2 times with similar results. **B)** Western blot analysis of scr and siPNUTS cells at 72 hrs after transfection. Bottom bar chart shows quantification of pRNAPII S5 relative to RNAPII (n=14). **C)** Western blot analysis of scr or siPNUTS transfected HeLa cells treated with thymidine for 2, 3, 4 and 6 hr. THZ1 was added at 2 hrs after thymidine to the indicated samples. *The bottom charts show quantification of pRNAPII S5 relative to CDK1, and pCHK1 S317 relative to CHK1. Values are shown for THZ1 and thymidine versus thymidine alone, for the respective siRNA oligos (n=4).

As the RNAPII CTD is the only known direct substrate of PNUTS-PP1 (Ciurciu et al., 2013, Lee et al., 2010), and RNAPII has a proposed role in ATR activation (Derheimer et al., 2007, Lindsey-Boltz & Sancar, 2007), we addressed whether dephosphorylation of RNAPII CTD is likely involved in the effects of PNUTS depletion on ATR signaling. We first verified that higher levels of pRNAPII S5 could be observed after depletion of PNUTS in HeLa cells (Fig 2B). We next added THZ1, a specific inhibitor of CDK7, the kinase mediating phosphorylation of RNAPII S5 (CTD) (Heidemann, Hintermair et al., 2013, Kwiatkowski, Zhang et al., 2014), to control siRNA versus PNUTS siRNA transfected cells during thymidine-induced replication stalling. To allow a robust activation of ATR signaling before inhibition of CDK7, thymidine was added two hours prior to THZ1. Remarkably, both pRNAPII S5 and pCHK1 S317 were reduced upon addition of THZ1 to cells transfected with control siRNA (Fig 2C, lanes 11-13), and both pRNAPII S5 and pCHK1 S317 remained high in PNUTS depleted cells (Fig 2C, lanes 14-16). This strongly indicates that pCHK1 S317 depends on pRNAPII S5. Of note, while the ATR inhibitor VE822 reduced pCHK1 S317 equally in both PNUTS depleted and cells transfected with control siRNA (Fig S2C), the CDK7 inhibitor THZ1 only reduced pCHK1 S317 in the control siRNA transfected cells (Fig 2C), thus ruling out the possibility that THZ1 should directly inhibit ATR kinase.

The finding that pRNAPII S5 levels remained high in PNUTS depleted cells after THZ1 treatment (Fig 2C) is consistent with a major role of PNUTS-PP1 in mediating the dephosphorylation of this residue (Fig 2C, compare lanes 14-16 with lanes 11-13). However, we observed that pRNAPII S7 and pRNAPII S2 also remained higher in PNUTS depleted cells under these conditions, though the effects appeared weaker than observed for pRNAPII S5 (Fig S3A). Therefore, PNUTS-PP1 may also participate in direct dephosphorylation of pRNAPII S2 and/or S7, or, dephosphorylation of S2 and S7 may depend upon S5. Furthermore, this shows that high ATR signaling correlates with RNAPII CTD phosphorylation in general, rather than with pRNAPII S5 specifically under these conditions.

To confirm the correlation between ATR signaling and RNAPII CTD phosphorylation, we added THZ1 to IR-treated cells. Similarly as observed during replication stress, pRNAPII S5 and pCHK1 S317/S345 were reduced after THZ1 in cells transfected with control siRNA (Fig S3B). And again, pRNAPII S5 and pCHK1 S317/S345 remained high in cells depleted for PNUTS (Fig S3B). An inhibitor of translation, cycloheximide, did not reduce pRNAPII S5 and pCHK1 S317/S345 after IR (Fig S3B), neither in control nor in PNUTS depleted cells, suggesting the effects of THZ1 on ATR signaling are independent of de novo protein production (via transcription and translation). To further explore the correlation between RNAPII CTD phosphorylation and ATR signaling, THZ1 was added prior to IR. As expected, pCHK1 S317 was suppressed by THZ1 in HeLa cells (Fig S3C). Similar effects were obtained with two other transcription inhibitors which also caused reduced RNAPII CTD phosphorylation, 5,6-Dichloro-1-β-D-ribofuranosylbenzimidazole (DRB) and triptolide, but not by cycloheximide (Fig S3C). Altogether these results support a link between RNAPII CTD phosphorylation and ATR signaling and indicate PNUTS-PP1 inhibits ATR activity by dephosphorylating pRNAPII CTD.

ATR is commonly known to be activated via replication stress, such as typically induced by hydroxyurea (HU). To compare ATR signaling in PNUTS depleted cells with ATR signaling after HU, we added different amounts of HU to non-transfected HeLa cells. HeLa cells treated with 60-100 μM HU for 24 hr showed equal or higher levels of replication stalling compared to PNUTS depleted cells 48 hr after siRNA depletion, as measured by lower uptake of the nucleoside analog EdU (Fig S4A). However, higher pCHK1 S317/S345 could still be observed in the PNUTS depleted cells (48 hr after siRNA transfection-Fig 3A), suggesting that ATR activity induced by depletion of PNUTS cannot simply be explained by replication stress. Notably, at 24 hr after siRNA transfection, there was no detectable difference in EdU uptake between control and siPNUTS transfected cells (Fig S4A), and therefore PNUTS depletion at 48 hours was comparable to 24 hours HU treatment.

**Figure 3.**
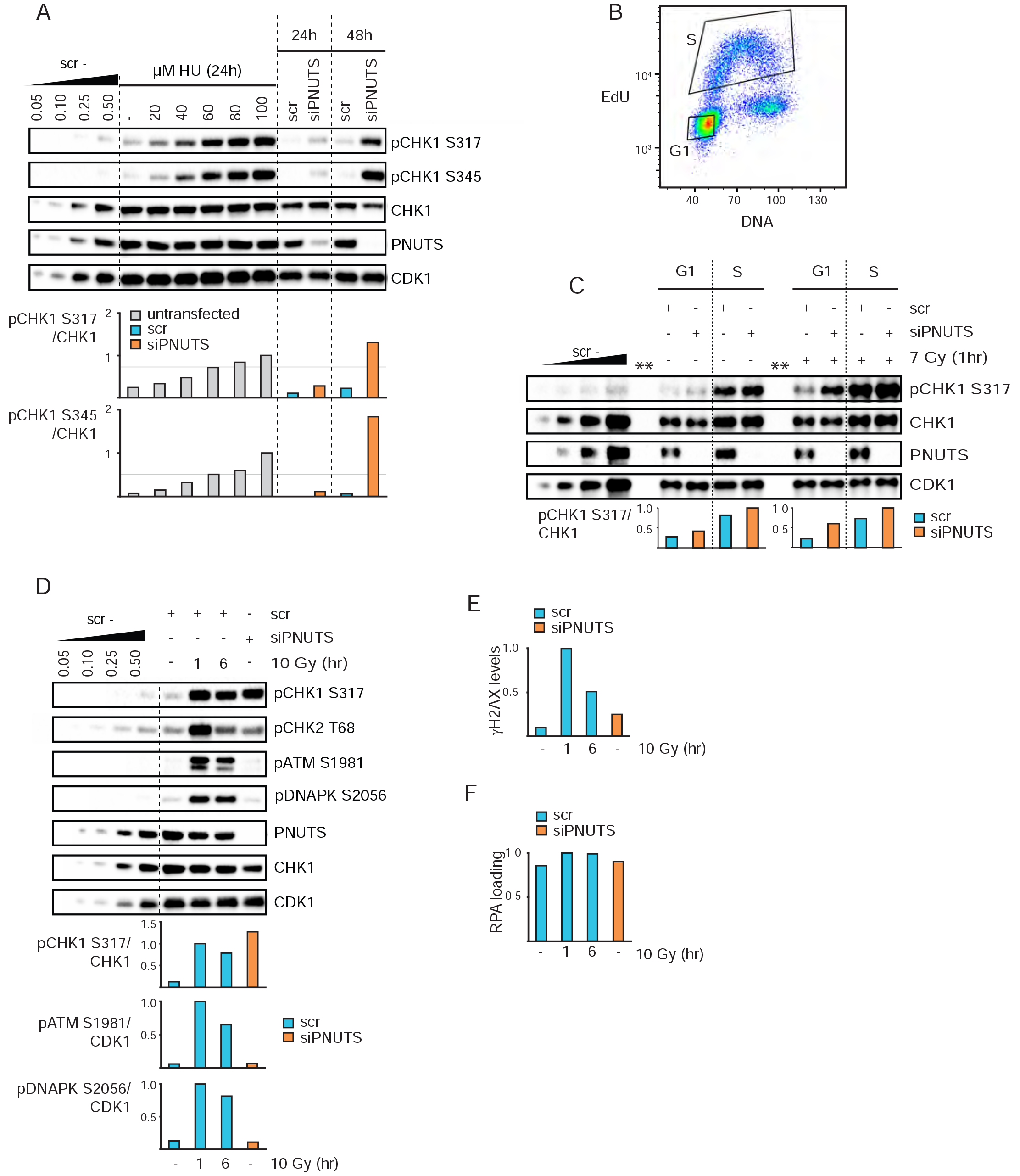
High ATR signaling after PNUTS depletion does not correlate with DNA damage markers or replication stress and can occur even in G1 phase. **A)** Western blot analysis of HeLa cells treated with indicated doses of HU for 24h and scr and siPNUTS transfected cells 24 and 48h after siRNA transfection (without HU). Bottom bar charts show quantifications of pCHK1 S317 and pCHK1 S345 relative to CHK1. The experiment was performed three times under similar conditions. Results are shown from one representative experiment. **B)** Cell sorting was performed by flow cytometry into G1 and S phases based on EdU incorporation and DNA content as indicated. **C)** Western blot analysis and quantifications of scr and siPNUTS transfected HeLa cells, sorted as in B and harvested at 48 hr after siRNA transfection, with and without IR (harvested at 1 hr after 7 Gy). Cells were irradiated immediately prior to addition of EdU and harvested at1 hr later. One representative image is shown, with ** indicating empty lanes. Quantifications were performed on images with different exposure times for the non-irradiated and irradiated samples (due to their different intensities), and normalized to the respective siPNUTS S phase sample. The experiment was performed three times, two at 72 hr and one at 48 hr with similar results. **D)** Western blot analysis of DNA damage markers for scr (without IR or 1 and 6 hr after 10 Gy) and siPNUTS transfected cells 48h after siRNA transfection. **E)** Bar chart showing median levels of yH2AX from flow cytometry analysis from cells harvested in parallel samples in the same experiment as in D. The analysis of parallel samples was performed in two independent experiments with similar results. **F)** Bar chart showing median levels of RPA loading from flow cytometry analysis of pre-extracted cells from the same experiment as in D. The experiment in F compared to D was performed three times with similar conditions and results.

We reasoned that phosphorylated RNAPII CTD might permit ATR activation even in the absence of replication e.g. in G1. To address this issue, cells in G1 and S phases of the cell cycle were sorted based on EdU incorporation and DNA content (Fig 3B). Remarkably, pCHK1 S317 was higher in both G1 and S phase after depletion of PNUTS, with and without IR (Fig 3C). To validate the purity of the G1 population following sorting, thymidine, which specifically targets S phase cells, was added for 30 min after EdU labeling (Fig S4B). Induction of pCHK1 S317 and presence of cyclin A could only be detected in the S phase population (Fig S4B), confirming pure populations. These results support that ATR signaling is increased even in G1 phase following PNUTS depletion.

ATR is also well known to be activated by DNA damage, such as occurring after IR (Jazayeri, Falck et al., 2006). We therefore next compared PNUTS depleted cells with IR-treated control siRNA transfected cells to address whether the high ATR activity after PNUTS depletion could correlate with DNA damage. Higher levels of DNA damage markers pATM S1981, pDNAPK S2056, pCHK2 T68 and γH2AX, but lower levels of pCHK1 S317, were observed in IR-treated control cells (1 and 6h after 10 Gy) compared to PNUTS depleted cells (Fig 3D,E). Furthermore, the lack of DNA damage signaling in PNUTS depleted cells was not caused by a reduced ability to activate ATM or DNAPK, as this occurred normally after IR (Fig S4C). The high ATR activity in PNUTS depleted cells is therefore not likely caused by DNA damage.

Moreover, we assessed RPA loading onto chromatin, which can occur both in response to DNA damage and replication stress. To measure RPA loading we used a flow cytometry based assay similar to one previously shown to detect end resection (Ferretti, Himmels et al., 2016). Higher RPA loading, but lower CHK1 S317 phosphorylation, could be observed in control cells after 10 Gy compared to PNUTS depleted cells (Fig 3F compared to 3D). This suggests a lack of correlation between RPA loading and ATR signaling after depletion of PNUTS. To further investigate this, we also codepleted PNUTS with RPA70, an essential component of the RPA complex (Iftode, Daniely et al., 1999). As expected, this resulted in strongly reduced pRPAS33 (Fig S4D,E). However, ATR-dependent CHK1 S345 phosphorylation was not reduced in cells codepleted for PNUTS and RPA70 compared to control cells or cells depleted for RPA70 alone (Fig S4D,E). The increased ATR signaling following depletion of PNUTS is therefore likely not due to increased amounts of ssDNA-RPA.

We also addressed the involvement of other known key upstream ATR activating proteins, namely TOPBP1 and ETAA1. Though pCHK1 S345 was reduced, ATR-dependent pRPA S33 was not reduced in cells co-depleted for TOPBP1 and PNUTS compared to cells depleted for PNUTS alone, in the absence or presence of IR (Fig 4A-C). Notably, enhanced pRPA S33 after IR in cells co-depleted for PNUTS and TOPBP1 was dependent on PNUTS as depletion of TOPBP1 alone did not cause increased pRPA S33 (Fig 4B,C). Conversely, upon co-depletion of PNUTS with ETAA1, pRPA S33 was reduced, but pCHK1 S345/S317 was not, compared to cells depleted of PNUTS alone (Fig 4D). Again the enhanced pCHK1 S317/S345 was dependent on PNUTS, as pCHKS317/S345 was not enhanced in cells depleted of ETAA1 alone compared to cells transfected with control siRNA (Fig 4D). Triple depletion of PNUTS, ETAA1 and TOPBP1 suppressed both pCHK1 S317/S345 and pRPA S33 (Fig 4D) in agreement with recent findings suggesting that TOPBP1 is required for pCHK1 S317/S345 and ETAA1 for pRPA S33 (Bass, Luzwick et al., 2016, Haahr, Hoffmann et al., 2016). Neither TOPBP1 nor ETAA1 therefore appear to be required for PNUTS dependent ATR signaling in general, but rather play essential downstream roles in the phosphorylations of CHK1 and RPA, respectively.

**Figure 4.**
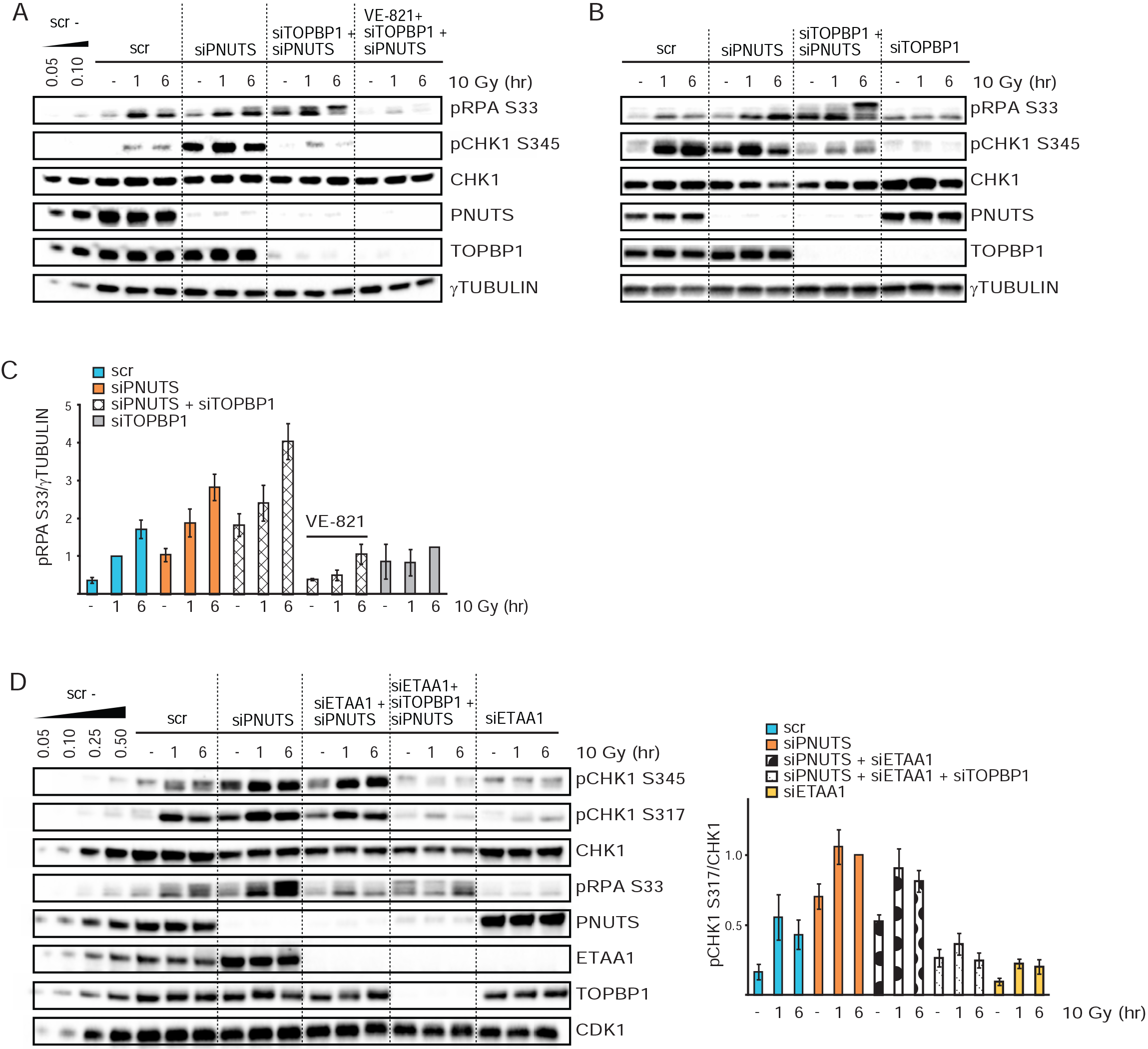
In cells depleted of PNUTS, TOPBP1 and ETAA1 mediate CHK1 and RPA phosphorylation, respectively. **A-C)** Western blot and quantifications (n=3) from cells transfected with scr, siPNUTS, and siRNA against TOPBP1 (siTOPBP1) harvested at 72 hrs after siRNA transfection and 1 and 6 hr after 10 Gy. VE-821 was added 30 min prior to 10 Gy. For siTOPBP1 10 Gy 6 hr sample error bar was emitted as experiment was performed two times. D) Western blot and quantifications (n=3) from cells transfected with scr, siPNUTS, siTOPBP1 and siRNA against ETAA1 (siETAA1) harvested at 48 hrs after siRNA transfection and 1 and 6 hr after 10 Gy.

To further characterize known ATR regulators following depletion of PNUTS, we compared their levels in cells transfected with PNUTS or control siRNA 24 or 48hr after siRNA transfection. Levels of ATR and ATRIP were not detectibly altered (Fig S5A). However, ETAA1, CLASPIN and TOPBP1 levels were increased in PNUTS depleted cells compared to control transfected cells, particularly at 48 hrs after siRNA transfection (Fig S5A). The co-depletions of PNUTS with ETAA1 or TOPBP1 nevertheless suggest that the ATR signaling can occur independently of either of these factors, though they are required for downstream phosphorylations (Fig 4). Also, as CLASPIN levels were downregulated, but pCHK1 S317 was higher after IR in PNUTS depleted cells relative to cells transfected with control siRNA (Fig S5B), this suggests CLASPIN is not either essential for enhanced ATR signaling upon PNUTS downregulation.

Our results showing a connection between RNAPII CTD phosphorylation and ATR signaling (Fig 2B,C and S3) suggest the CTD may be acting as a signaling platform for ATR activity. We therefore searched for factors that might participate in signaling from phosphorylated RNAPII CTD towards ATR. In the literature, we identified three proteins, BRCA1, PRP19 and CDC73, that associate with hyperphosphorylated RNAPII and have been linked to ATR (David, Boyne et al., 2011, Krum, Miranda et al., 2003, Marechal, Li et al., 2014, Phatnani, Jones et al., 2004, Poli, Gerhold et al., 2016, Turner, Aprelikova et al., 2004). We found that co-depletion of BRCA1 or PRP19 with PNUTS did not reduce the high ATR signaling (data not shown). However, co-depletion of CDC73 with PNUTS reduced both pCHK1 S317/S345 and pRPA S33, but not pRNAPII S5, in the presence or absence of IR (Fig 5A). The reduction in pCHK1 S345 phosphorylation after co-depletion was observed with several siRNA oligos against CDC73 (four out of five) (Fig S5C). Furthermore, expression of siRNA resistant Flag-CDC73 partially rescued the effects on pCHK1 S317/S345 and pRPAS33 downregulation after co-depletion of CDC73 with PNUTS (Fig 5A), excluding siRNA off-target effects. These results suggest that CDC73 and PNUTS are acting in the same pathway for ATR activation and are consistent with a role for CDC73 in signaling from phosphorylated RNAPII CTD to ATR.

**Figure 5.**
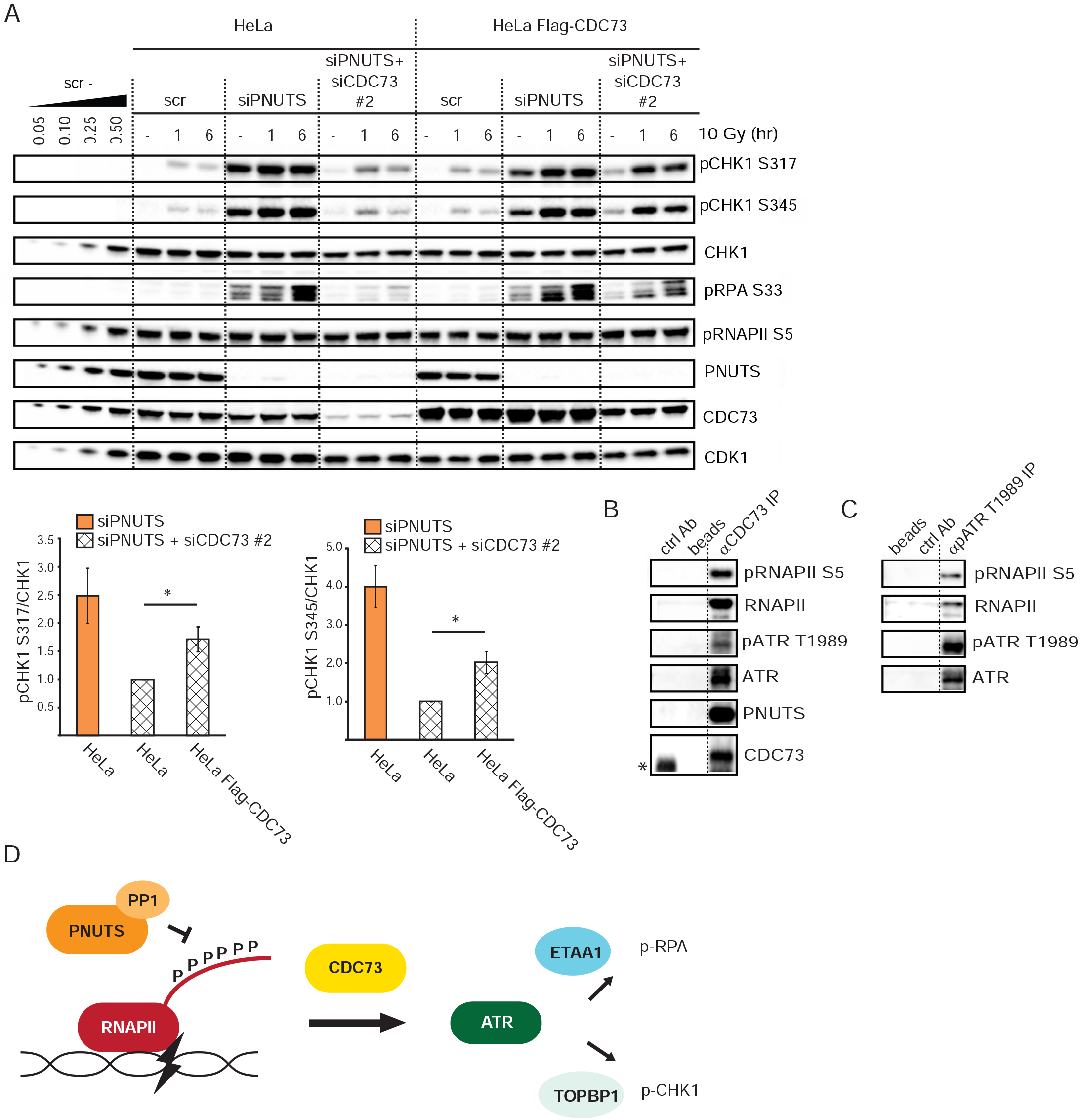
CDC73 interacts with ATR and RNAPII and is required for the high ATR signaling after PNUTS depletion. **A)** Western blot analysis and quantifications of scr, siPNUTS or CDC73 siRNA (siCDC73 #2) transfected HeLa cells or HeLa cells stably expressing siRNA-resistant Flag-CDC73 treated with IR (10Gy) as indicated. Bottom bar charts show quantification of pCHK1 S345 and pCHK1317 versus CHK1 levels at 6 hours after 10 Gy (n=3). *indicates p<0.05 by two-tailed T-test. **B)** Western blot analysis of immunoprecipitations from HeLa cell lysates, using a control antibody (ctrl Ab), no antibody (beads) or anti-CDC73 antibodies (aCDC73 IP). *Indicates IgG band from the control antibody, which migrated slightly faster than CDC73 in the western blot. **C)** Western blot analysis of immunoprecipitations as in B, but using anti-pATR T1989 antibodies(apATR T1989 IP). **D)** Model for how PNUTS-PP1 may suppress ATR signaling. Based on our results, we propose a pathway where RNAPII-CTD signals to ATR via CDC73. PNUTS-PP1 counteracts this by dephosphorylating the CTD. Furthermore, our results indicate that TOPBP1 and ETAA1 can direct the ATR activity towards pCHK1 S317/345 and pRPA S33, respectively.

CDC73 interacts genetically with the ATR homologue Mec1 in *Saccharomyces cerevisiae*, and a physical interaction has been proposed but not previously shown (Poli et al., 2016). To address whether CDC73 physically interacts with ATR and RNAPII, we performed co-immunoprecipitation (co-IP) experiments of endogenous proteins in HeLa cells. Indeed, co-IPs using a CDC73 antibody pulled down RNAPII, pRNAPII S5, ATR and pATR T1989 (Fig 5B). As pATR T1989 is thought to be an autophosphorylation site (Liu, Shiotani et al., 2011), this indicates that catalytically active ATR associates with CDC73. Interestingly, PNUTS and PP1 were also detected in the CDC73 co-IPs (Fig 5B and S5D). We verified that the immunoprecipitations were specific by using lysates from cells depleted of CDC73, which pulled down less ATR and RNAPII (Fig S5D). Notably, RNAPII is a well known interactor of CDC73 ((Rozenblatt-Rosen, Hughes et al., 2005) and reviewed in (Van Oss, Cucinotta et al., 2017)). Furthermore, the depletion of CDC73 was only partial and significant amounts of CDC73 were present in the co-IPs from cells transfected with CDC73 siRNA (Fig S5D, CDC73-high exposure), which may explain the residual ATR and RNAPII pulled down under these conditions. To address whether ATR and pRNAPII S5 could physically associate, we performed ATR co-IPs. To enrich for active ATR in these experiments, we used pATR T1989 antibodies. This efficiently pulled down ATR and faint bands corresponding to pRNAPII S5 and RNAPII could also be detected, suggesting an interaction in live cells (Fig 5C). All the co-IPs were performed after treatment with the endonuclease benzonase, strongly suggesting that the interactions were not mediated by DNA. Altogether these results support a role for phosphorylated RNAPII and CDC73 in the high ATR activity after PNUTS depletion.

## Discussion

ATR kinase plays a central role in signaling after DNA damage and replication stress. Here we show for the first time that the RNAPII phosphatase PNUTS-PP1 suppresses ATR signaling. Our results suggest that ATR signaling is restrained by PNUTS-PP1 mediated dephosphorylation of RNAPII CTD, and thus support a role for RNAPII in ATR signaling. Furthermore, we have found that a well-known RNAPII binding factor, CDC73, is required for the high ATR signaling in PNUTS depleted cells. Moreover, our results support recent findings that TOPBP1 and ETAA1 may direct ATR activity towards different substrates. Altogether, based on these results we propose a new model for ATR signaling via RNAPII (Fig 5D).

Notably, the high ATR signaling in PNUTS depleted cells does not correlate with DNA damage or replication stress and can occur even in G1 phase of the cell cycle (Fig 3). Signaling to ATR by pRNAPII CTD may therefore be a more general event that simply responds to RNAPII stalling, in line with previous reports showing that perturbation of transcription can induce ATR activation in the absence of DNA damage and prior to detection of replication-stress (Derheimer et al., 2007, Kotsantis, Silva et al., 2016). The *Saccharomyces cerevisae* ATR homologue Mec1 was shown to promote removal of RNAPII at sites of transcription-replication conflict (Poli et al., 2016). Viewed in light of our results, ATR activity at sites of stalled transcription might thus promote removal of RNAPII, even in the absence of replication. Removal of stalled RNAPII is likely important also outside of S phase, because RNAPII could create an obstacle for further transcription in a region which might e.g. contain an essential-or tumor suppressor gene. In agreement with prolonged RNAPII stalling being detrimental to the cell, it has been shown to be a strong signal for apoptosis (Ljungman & Zhang, 1996).

We found that pRNAPII CTD was required for, but did not strictly correlate with, ATR signaling (e.g. Fig S3C - compare lanes 1 and 2, pCHK1 S317 vs pRNAPII S5). However, as RNAPII CTD phosphorylation occurs during the normal transcription cycle, where e.g. pRNAPII S5 is required for 5’ mRNA capping (Egloff et al., 2012), a strict linear correlation may be unlikely as it would imply ATR activation merely as a consequence of normal transcription. We suggest that the level of pRNAPII CTD at individual sites may determine the degree of ATR activation by RNAPII. Perhaps ATR is activated only when pRNAPII CTD reaches an abnormal threshold level at an unexpected site. Supporting this, RNAPII CTD phosphorylation is often associated with RNAPII stalling, and can occur within the entire transcribed region (Alexander et al., 2010, Boehm et al., 2003, Munoz, Perez Santangelo et al., 2009). Furthermore, other post-translation modifications on the CTD (Harlen & Churchman, 2017) may contribute to fine-tune ATR activity via RNAPII.

Of note, in the alternative splicing response to UV, pRNAPII CTD was proposed to occur downstream of ATR activation, and ATR activation to occur independently of transcription in HaCaT cells (Munoz, Nieto Moreno et al., 2017). These results may appear to be contradictory to ours. However, we did not detect any reduction in pRNAPII S5 after ATR inhibitor during replication stress in HeLa cells (Fig S2C) suggesting ATR is not always upstream of pRNAPII CTD. Furthermore, the differing results may be explained by the existence of several pathways for ATR activation acting in parallel, e.g. via RNAPII, via ssDNA-RPA, and via unknown pathways. The contribution from each pathway is likely to vary between cell types and with different stresses.

Remarkably, following depletion of PNUTS ATR signaling did not correlate with RPA loading and was not reduced by RPA70 co-depletion (Fig 3D,F and S4D,E). These results are in agreement with previous studies showing that RPA loading and ATR activation do not always correlate (Dodson, Shi et al., 2004, Kousholt, Fugger et al., 2012). However, in our co-depletion experiments, we cannot exclude that very small amounts of ssDNA-RPA (less than 5% of RPA70 was remaining after siRNA depletion (Fig S4D)) may contribute to the ATR activity. Indeed, recently, ATR activation was shown to occur at centromeres in mitosis, by a proposed mechanism involving R-loops and ssDNA-RPA (Kabeche, Nguyen et al., 2017). The authors show the dependency on R-loops by overexpression of RNAse H which reduced R-loops and pCHK1 S317 (Kabeche et al., 2017). As phosphorylated RNAPII CTD can cause formation of R-loops (Kaneko, Chu et al., 2007), which can subsequently lead to small amounts of ssDNA-RPA (Nguyen, Yadav et al., 2017), it is thus tempting to speculate that depletion of PNUTS may cause small amounts of ssDNA-RPA via R-loops, and thereby enhance ATR signaling. On the other hand, there is an intimate connection between stalled RNAPII and R-loops (Santos-Pereira & Aguilera, 2015). Interestingly, it was recently shown that overexpression of RNAse H can also cause release of stalled RNAPII, suggesting that R loops can promote RNAPII stalling (Sridhara, Carvalho et al., 2017). Another speculation might therefore be that R-loops could activate ATR by leading to stalling of RNAPII and subsequent RNAPII CTD phosphorylation.

Our results point to a new role for CDC73 upstream of ATR activation. CDC73 is a component of the PAF1 complex, including PAF1, CTR9, LEO1, RTF1 and WDR61, involved in all stages in RNAPII transcription (Van Oss et al., 2017). However, the PAF1 complex does not appear to be essential for transcription as depletion of e.g. CDC73 was found to both up and down-regulate mRNA expression (Rozenblatt-Rosen, Nagaike et al., 2009). In *Saccharomyces cerevisae* the PAF1 complex was required for removal of RNAPII during collisions of transcription and replication (Poli et al., 2016). Leo1 is a target of Mec1, and thus Mec1 has been thought to primarily have an upstream role in triggering the removal of RNAPII via phosphorylation of Leo1 and the Ino80 chromatin remodeling complex (Hustedt et al., 2015, Poli et al., 2016). However, as our results point to a role for CDC73 in ATR activation, this may suggest the existence of a feedback loop, whereby stalling of RNAPII in head-on collisions between the transcription and replication machineries leads to activation of ATR via CDC73, and the resultant ATR activity leads to removal of RNAPII. Interestingly, our results fit well with the observation that head-on collisions, which cause disruption of RNA transcription, lead to ATR activity, whereas co-directional collisions, which do not disrupt RNA transcription, do not (Hamperl, Bocek et al., 2017).

CDC73 is also a well known tumor suppressor gene. It is currently not clear how CDC73 acts as a tumor suppressor, though roles in Wnt signaling, regulation of P53 and CYCLIN D levels and homologous recombination repair have been suggested (Herr, Lundin et al., 2015, Jo, Chung et al., 2014, Mosimann, Hausmann et al., 2006, Woodard, Lin et al., 2005). ATR activity protects genome integrity by stabilizing stalled forks during replication stress and promoting DNA repair and checkpoint activation (Yazinski & Zou, 2016). However, in addition ATR activity can promote apoptosis in non-cycling cells, which are the majority of cells in humans (Kemp & Sancar, 2016). Therefore, CDC73 could potentially protect against cancer by promoting RNAPII-mediated ATR activity leading to cell death in non-cycling cells with DNA damage. Consistent with this interpretation, PNUTS, which counteracts CDC73 in ATR activation, is a putative proto-oncogene (Kavela, Shinde et al., 2013).

## Materials and methods

### Cell culture and treatments

Human epithelial HeLa and human osteosarcoma U2OS cells were grown in Dulbecco’s Modified Eagle Medium (DMEM) containing 10% fetal calf serum (Life Technologies). The cell lines were authenticated by short tandem repeat profiling using Powerplex 16 (Promega) and regularly tested for mycoplasma contamination. HeLa BAC cells stably expressing EGFP mouse pnuts were a generous gift from the laboratory of Tony Hyman (http://hymanlab.mpi-cbg.de/bac_viewer/search.action). To generate the flag-CDC73 cell lines, CDC73 (Addgene plasmid # 11048) was amplified using the primers aggctttaaag-gaaccaattcagtcgactgGAATTCGGATCCACCA (Cdc73 entry fwd) and aagaaagctgggtctagatatctcgagtgcTCAGAATCTCAAGTGCG (Cdc73 entry rev) and cloned into BamH1-Not1 cut pENTR1A using Gibson cloning (NEB E5510S). To generate the siRNA-resistant constructs, silent mutations were introduced in the siRNA target site using the Quick Change Lightning kit (Agilent 210518). The mutagenic primers were: CATCAGATGAAAAGAAGAAGCAGGGA-TGCCAGAGGGAAAATGAAACTCTAATACA and TGTATTAGAGTTTCATTTTCC-CTCTGGCATCCCTGCTTCTTCTTTTCATCTGATG. The construct was cloned into the lentiviral expression vector pCDH-eF1-GW-IRES-puro by Gateway cloning (Thermo-Fisher Scientific 11791020). HeLa cells were transduced and cells carrying the transgene were selected with 0,5 μg/ml puromycin.

Cells were irradiated in a Faxitron x-ray machine (160 kV, 6.3 mA, 1 Gy/min). Thymidine (Sigma-Aldrich) was used at 2 mM, ATR-inhibitors VE-821 (Axon Medcem) and VE-822 (Selleck Biochem) at 10 μM and 1 μM respectively, CDK7-inhibitor THZ1 (ApexBio) at 1 μM, CDK9-inhibitor DRB (Sigma-Aldrich) at 100 μM, XPB-inhibitor triptolide (Sigma-Aldrich) at 1 μM and translational inhibitor cycloheximide (Sigma-Aldrich) at 10 μg/ml.

### siRNA and DNA transfections

Wildtype and RAXA (mutated in the ‘RVXF’ (^398^SVTW^401^) motif: V399A, W401A) fullength EGFP PNUTS DNA constructs containing 14 silent mutations in the domains targeted by siPNUTS (#1 and #2) were synthesized by Geneart and cloned into pGLAP3. Sequences of siRNA oligonucleotides can be found in supplemental table 1. siRNA was transfected using Oligofectamine or RNAimax (Life technologies), and plasmid DNA with Fugene HD (Promega) or Attractene (Qiagen). Experiments were performed 65-72 hours after siRNA transfection unless otherwise stated.

### Western blotting and antibodies

For quantitative western blotting, cells were resuspended in ice-cold TX-100 buffer (100 mM NaCl, 50 mM Tris pH 7.5, 2 mM MgCl2, 0.5 % TX-100) containing 100 U/ml Benzonase (Sigma-Aldrich). After 1 hr incubation on ice, Lane Marker Reducing Sample Buffer (Pierce Biotechnologies) was added and samples were boiled (95°C, 5 min). Criterion TGX gels (BioRad) and nitrocellulose membranes (BioRad) were used for separation and transfer respectively. Antibodies were: anti-PNUTS (BD Biosciences), anti-phosphoCHK1 Ser317, anti-phosphoCHK1 Ser345, anti-phosphoATM Ser1981, anti-phosphoCHK2 Thr68, anti-ATR (all from Cell Signaling Technology), anti-CHK1 and anti-CHK2 (both (Sorensen, Syljuasen et al., 2003)), anti-γTUBULIN (GTU-88, Sigma), anti-CDK1 (sc-54), anti-RNAPII (F-12), anti-MCM7 (DCS-141) (all from Santa Cruz Biotechnology), anti-phospho RPA32 Ser33, anti-phosphoDNAPK S2056 (both from Abcam), anti-phospho RNAPII S5 (3E8), anti-phospho RNAPII S7 (4E12), anti-phospho RNAPII S2 (3E10) (all from Millipore), anti-CDC73 (Bethyl), anti-phosphoATR Thr1989 (GeneTex) and anti-ETAA1 (Haahr et al., 2016). Peroxidase-conjugated secondary antibodies were from Jackson Immunoresearch. Blots were imaged in a Chemidoc MP (BioRad) using chemiluminescence substrates (Supersignal west pico, dura or femto; Thermo Scientific). Quantifications were performed and images processed in Image Lab 4.1 (BioRad) software. Linearity of detection was verified by including a dilution series of one of the samples (see e.g. Figure 1B) and excluding saturated signals. The resulting standard curve allowed accurate quantification. To blot for total protein after detection of a phosphorylated protein, membranes were stripped using ReBlot Plus Mild Antibody Stripping Solution (Millipore).

### Cell sorting and flow cytometry

For cell sorting and flow cytometry with EdU labeling, cells were labeled for 1 hr with 2 μM EdU and fixed in 70% ethanol. EdU was labeled with the Click-iT Plus EdU Alexa Fluor 488 Flow Cytometry Assay Kit (Thermo Fisher), and DNA with FxCycle Far Red. Cells were sorted with a BD FacsAria Cell Sorter (BD Biosciences) using FlowJo software. Sorted cells were analyzed by western blotting as above. For flow cytometry of RPA loading, cells were pre-extracted, fixed and labeled as in (Haland, Boye et al., 2015) using anti-RPA70 antibodies (Cell Signaling). For flow cytometry of γH2AX, samples were fixed and labeled as in (Hauge, Naucke et al., 2017). For analysis, a LSRII flow cytometer (BD Biosciences) was used with Diva or FlowJo software.

### Immunoprecipitation experiments

For immunoprecipitations, cells were lyzed in TX-100 buffer (see under western blotting) containing 100 U/ml Benzonase (Sigma-Aldrich). Lysates were precleared and anti-CDC73 (Bethyl) or anti-phosphoATR Thr1989 (GeneTex) or anti-pCHK2 T68 (used as control antibody, from Cell Signaling) were added. Dynabeads (protein G; Life technologies) were used to isolate antibody-bound complexes.

### Statistics

All experiments, except when otherwise stated, were performed three times or more. Error bars represent standard error of mean (SEM).

## Acknowledgements

We thank Libor Macurek, Sebastian Patzke, Jiri Lukas and Claus Storgaard Sørensen for useful discussions and Niels Mailand for providing the ETAA1 antibody. This work was supported by grants from the Norwegian Cancer Society, the South-Eastern Norway Health Authorities and the EEA Czech-Norwegian Research Programme (Norwegian Financial Mechanism 2009-2014 and the Ministry of Education, Youth and Sports under Project Contract no MSMT-22477/2014 (7F14061)).

## Author contributions

HBL performed most of the experiments. HBL and RGS conceived the project and analyzed the results. LES and GER performed experiments and contributed to the analysis. BG constructed HeLa cells expressing siRNA resistant CDC73. LTM contributed conceptually with regards to PP1 and to the development of cells expressing GFP-PNUTS. HBL and RGS wrote the paper. All the authors contributed to revision of the paper.

## Supplementary figure legends

**Figure S1.**
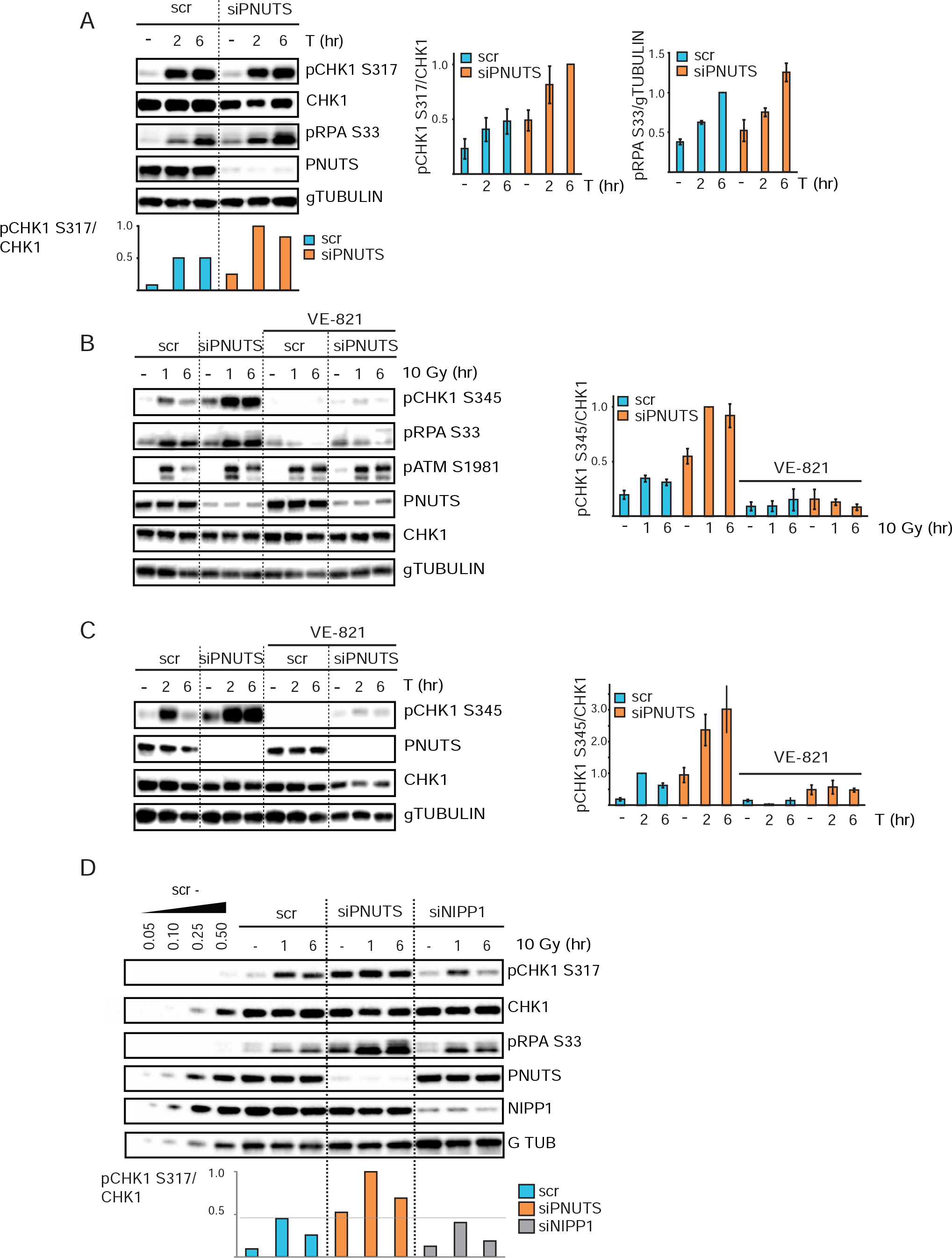
**A)** Western blot analysis and quantifications (n=3) of scr and siPNUTS transfected U2OS cells at 2 or 6 hr after addition of thymidine. Bar chart under the western blot show results from the same experiment. Bar charts to the right show quantification from 3 independent experiments. **B)** Western blot analysis and quantifications (n=3) of scr and siPNUTS transfected HeLa cells at 1 or 6 hr after 10 Gy. VE-821 was added 30 min prior to IR. **C)** Western blot analysis and quantifications (n=3) of siPNUTS HeLa cells at 2 or 6 hr after thymidine. VE-821 was added 30 min prior to thymidine. **D)** scr, siPNUTS cells or cells transfected with siRNA against NIPP1 (siNIPP1) at 1 or 6 hr after 10 Gy. Experiment was performed two times with similar results.

**Figure S2.**
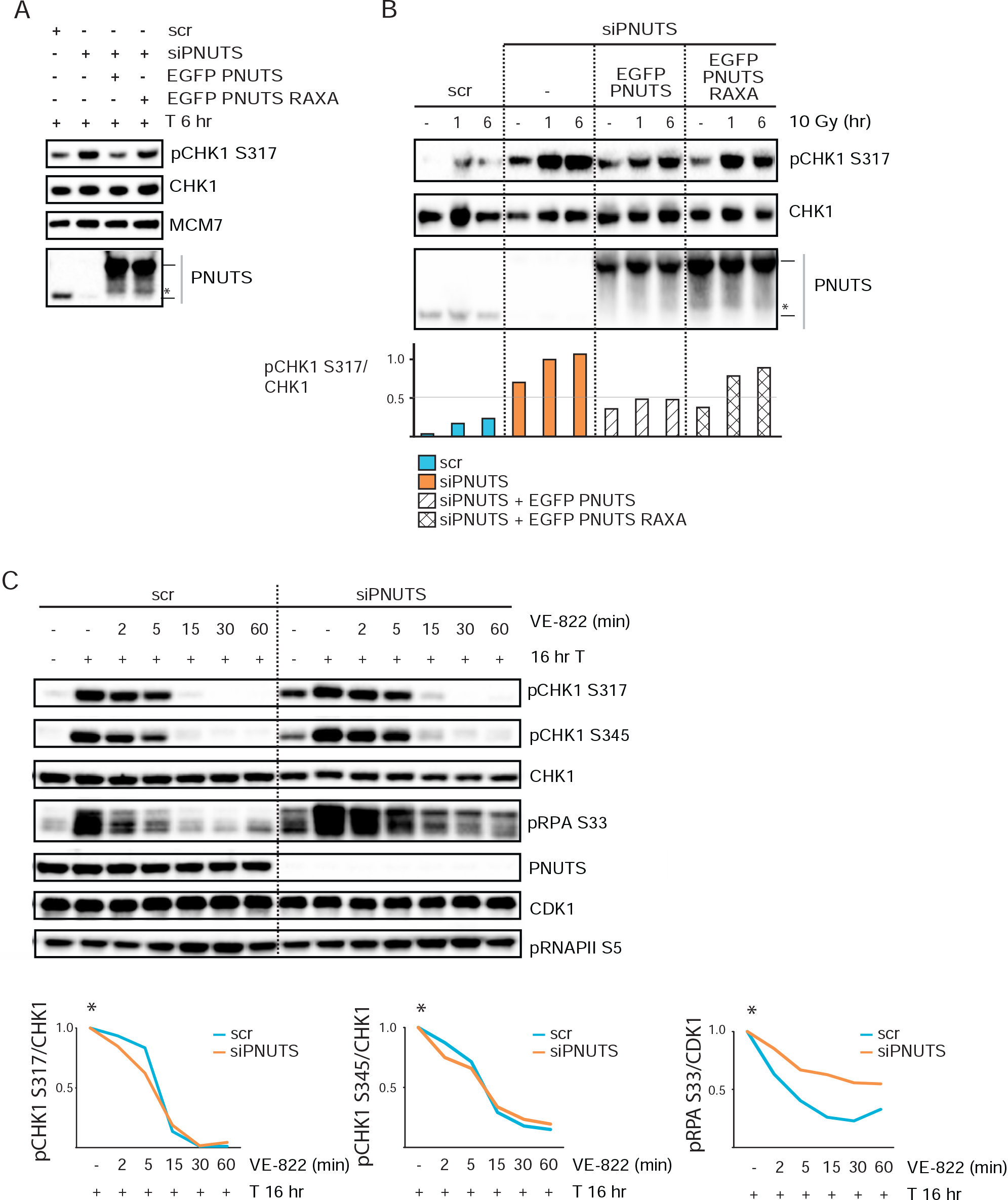
**A)** Representative western blot from experiment as in 1D, 6 hr after thymidine. **B)** Western blot analysis and quantifications from experiment performed as in 1D). **C)** Western blot analysis of scr or siPNUTS transfected cells without or with thymidine (16hr T). VE822 was added for 2, 5, 15, 30, or 60 min to indicated samples 16hr after addition of T. *Charts show values of pCHK1 S317/CHK1 in cells treated with T and VE822 vs T 16 hr alone, for respective siRNA oligos. Experiment was performed two times with similar results.

**Figure S3.**
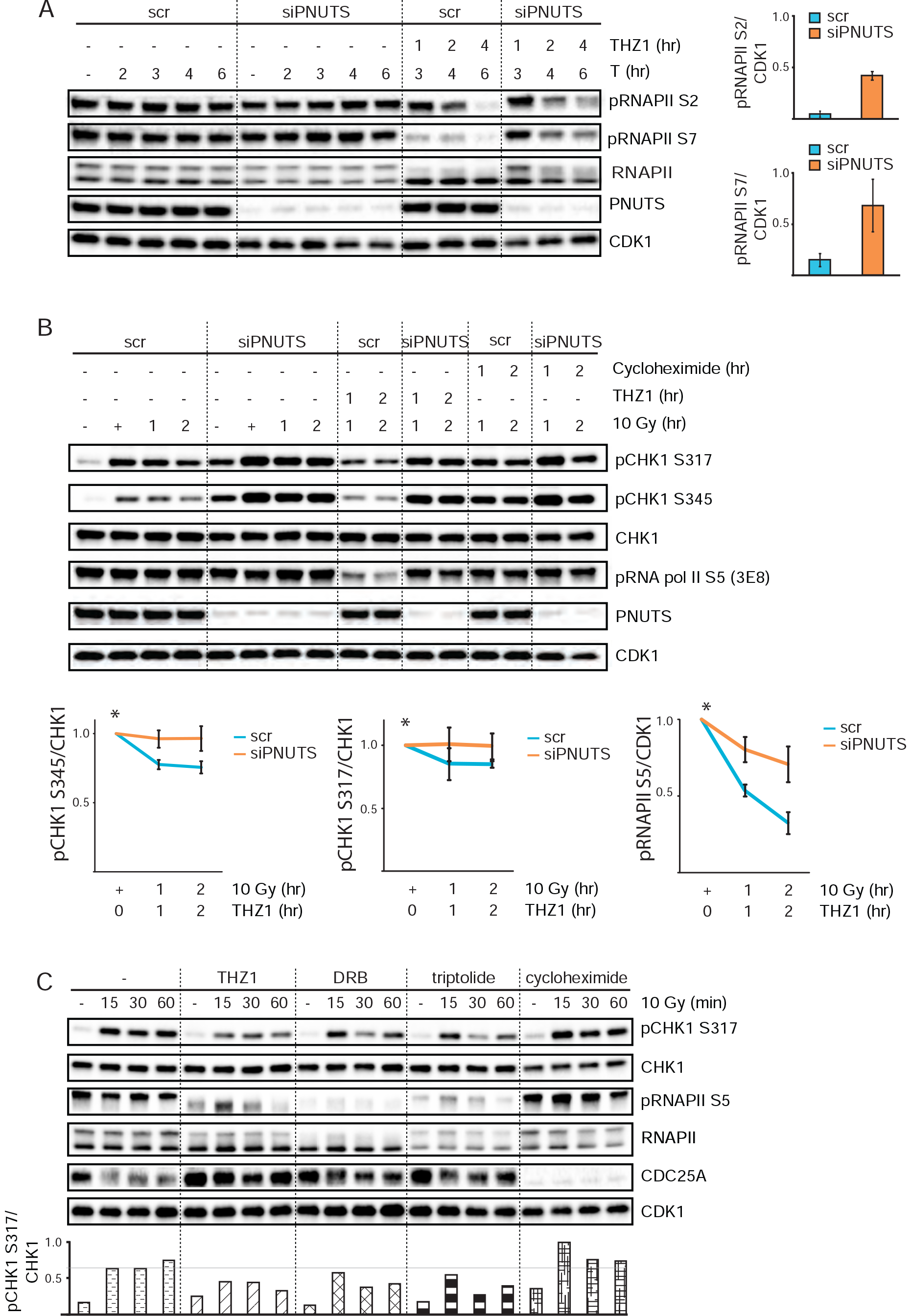
**A)** Representative western blot and quantification of experiment performed as in 2C. **B)** Western blot analysis of scr or siPNUTS transfected cells without or with IR (10 Gy, 1 or 2 hrs). THZ1 or cycloheximide was added 10 min after IR to the indicated samples. Samples with or without THZ1 were collected together to allow direct comparison. The bottom charts show quantification of pRNAPII S5 relative to CDK1 and pCHK1 S317/S345 relative to CHK1 levels (n=3). *Values are shown for THZ1 + 10 Gy vs 10 Gy alone for the respective siRNA oligos. **C)** Representative western blot and resulting quantification from HeLa cells with or without transcriptional inhibitors THZ1, DRB, triptolide or translational inhibitor cycloheximide. Inhibitors were added to cells 60 min prior to 10 Gy and samples were harvested at 15, 30 and 60 min. CDC25A levels verify effects of cycloheximide on a short-lived protein. Quantifications beneath western blot shows results from the same experiment. The experiment was performed three times under resembling conditions with similar results.

**Figure S4.**
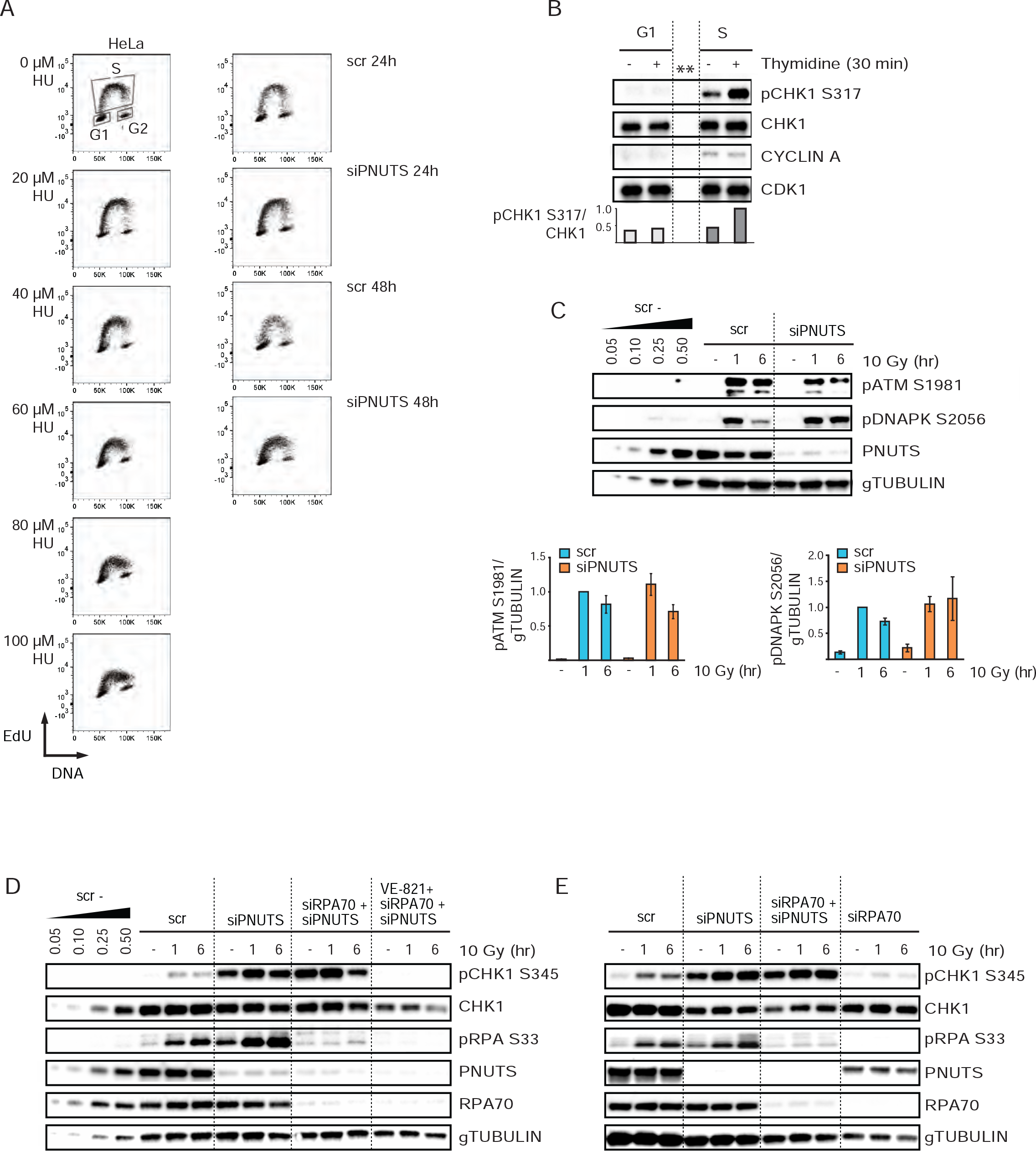
**A)** Charts showing EdU versus DNA content from parallel samples harvested within the same experiment as in 3A. **B)** Western blot and quantifications from cell cycle sorting experiment as in 3B. Cells were labeled with EdU for 1 hr, followed by 30 min incubation with thymidine, prior to harvest. **C**) Western blot analysis and quantifications of scr or siPNUTS transfected cells without or with IR (10 Gy, 1 or 6 hrs)). (n=7 for pATM S1981, and n=5 for pDNAPK S2056). **D) and E)** Representative western blots of cells transfected with scr, siPNUTS, and siRNA against RPA70 (siRPA70) harvested at 72 hrs after siRNA transfection and 1 and 6 hr after 10 Gy. VE-821 was added 30 min prior to 10
Gy. (n=3).

**Figure S5.**
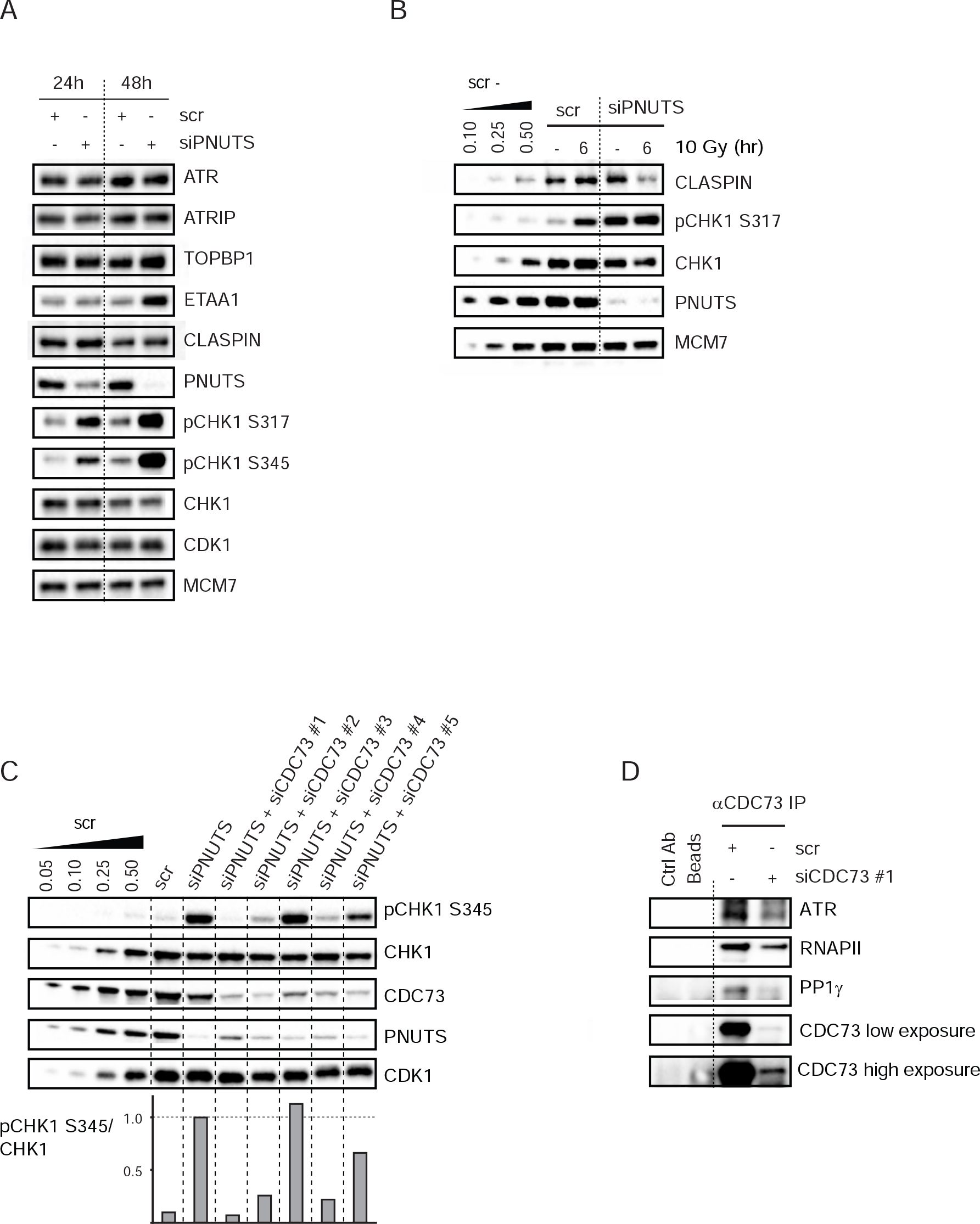
**A)** Western blot analysis of scr or siPNUTS transfected cells at 24 and 48hr after siRNA transfection. **B)** Western blot analysis of scr or siPNUTS transfected cells at72 hr after siRNA transfection, without and with IR (10 Gy, 6h). **C)** Western blot and quantifications from representative experiment of cells transfected with scr, siPNUTS, and five different oligonucleotides against CDC73, siCDC73 #1, #2, #3, #4 and #5 harvested 48 hr after siRNA transfection. (n=3). **D)** Western blot from immunoprecipitation experiment performed as in 5B on lysates from cells transfected with scr or siCDC73 #1, at 72 hr after siRNA transfection.

**Table S1.**
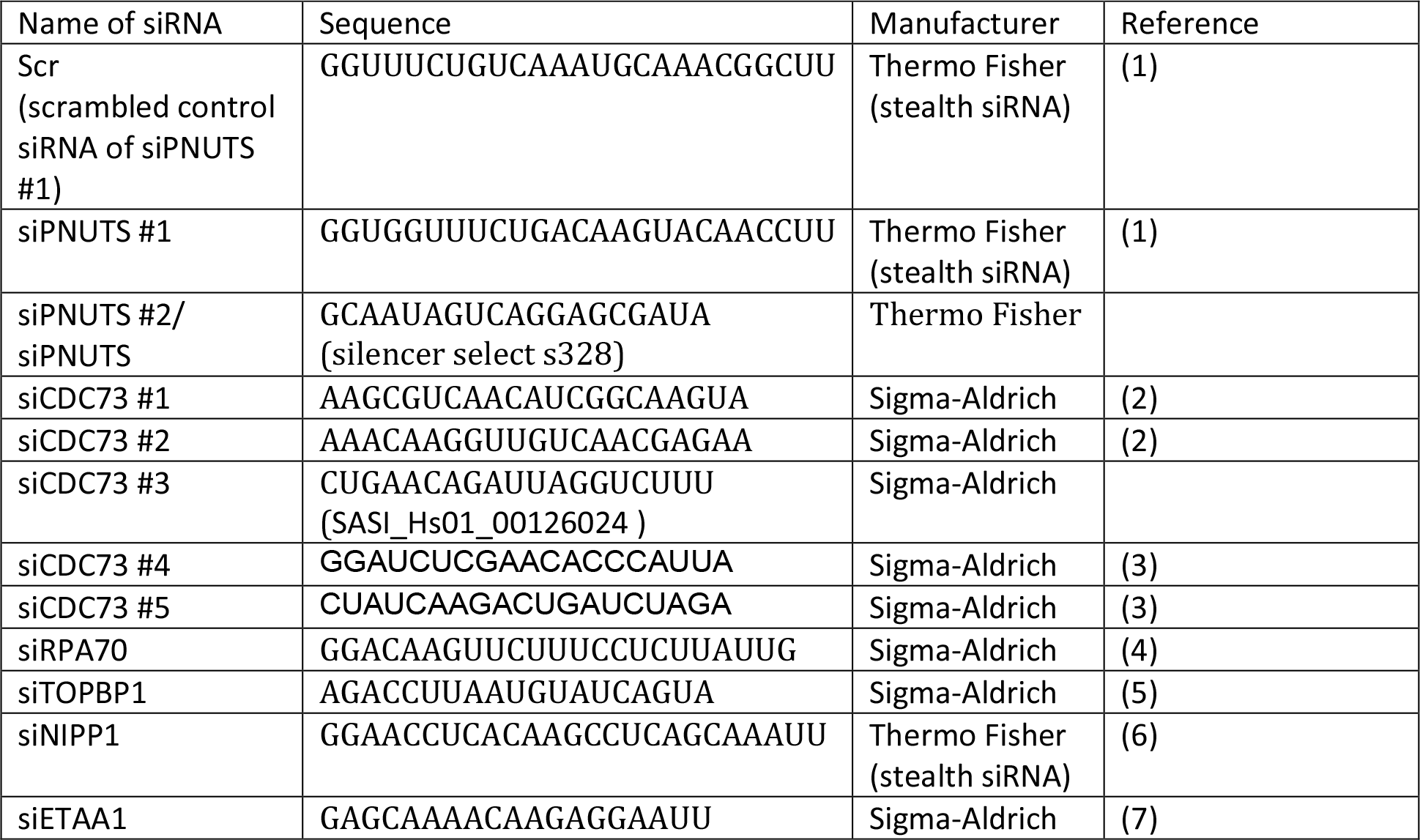
siRNA oligonucleotide sequences

